# Dysregulation of FicD AMPylation causes diabetes by disrupting pancreatic endocrine homeostasis

**DOI:** 10.1101/2024.10.25.620287

**Authors:** Amanda K. Casey, Nathan M. Stewart, Naqi Zaidi, Hillery F. Gray, Hazel A. Fields, Masahiro Sakurai, Carlos A. Pinzon-Arteaga, Bret M. Evers, Jun Wu, Kim Orth

## Abstract

Bi-functional enzyme FicD regulates the endoplasmic reticulum chaperone BiP using AMPylation and deAMPylation during ER homeostasis and stress, respectively. Human FicD with an arginine-to-serine mutation disrupts FicD deAMPylation activity resulting in severe neonatal diabetes. We generated the *FicD*^*R371S*^ mutation in mice to create a pre-clinical murine model for neonatal diabetes. We observed elevated BiP AMPylation levels across multiple tissues and signature markers for diabetes including glucose intolerance and reduced serum insulin levels. While the pancreas of *FicD*^*R371S*^ mice appeared normal at birth, adult *FicD*^*R371S*^ mice displayed disturbed pancreatic islet organization that progressed with age. *FicD*^*R371S*^ mice provide a preclinical mouse model for the study of UPR associated diabetes and demonstrate the essentiality of FicD for tissue resilience.

## Main Text

Insulin synthesis and secretion is a highly complex process involving multiple steps in protein processing and trafficking. Throughout the lifetime of an organism, β-cells must constantly adapt to fluctuating insulin demands. Disruption of endoplasmic reticulum (ER) homeostasis and dysregulation of the unfolded protein response (UPR) are key contributors to β-cell dysfunction and death in type I diabetes (*1-3*). While β-cells are generally resistant to transient ER stress, prolonged or chronic ER stress leads to impaired function and a loss of β-cell identity (*3, 4*). Mutations and disruptions in the UPR pathway have been implicated in both type I and syndromic diabetes (*4, 5*).

FicD is an ER-resident enzyme responsible for the reversible AMPylation of BiP in response to changes in ER stress (*6*). Recently, a recessive, disease-causing mutation in the human FicD gene (*hFicD*^*R371S*^) was identified as a cause of infancy-onset diabetes (*7*). The mutation disrupts FicD’s enzymatic activity, specifically impairing its ability to deAMPylate BiP in the ER (*7*). However, how this mutation impacts UPR regulation across different cell types and leads to neonatal diabetes remains unclear. In this study, we investigate the effects of the *FicD*^*R371S*^ mutation on pancreatic islet function by developing a pre-clinical murine model carrying the *FicD*^*R371S*^ mutation.

### Generation of a neonatal diabetic *FicD*^*R371S*^ mouse model

We utilized CRISPR-Cas9 homology-directed repair in C57Bl/6 zygotes to introduce the *FicD*^*R371S*^ mutation in mice (**Fig. 1, Fig. S1**). From these founders, we established a *FicD*^*+/R371S*^ mouse line, which was used for subsequent experiments. *FicD*^*R371S/R371S*^ mice were born at expected Mendelian ratios, appeared normal in body size, and displayed no overt health issues compared to their *FicD*^*+/+*^ and *FicD*^*+/R371S*^ littermates (**Fig. 1B-C**). To assess whether the *FicD*^*R371S*^ mutation recapitulates the diabetic phenotypes in human patients, we measured blood glucose and serum insulin levels in *FicD*^*R371S/R371S*^ mice and their sibling controls fed on a normal diet. By 5 weeks of age, *FicD*^*R371S/R371S*^ mice exhibited elevated blood glucose and reduced serum insulin levels (**Fig. S2**). Glucose tolerance tests (GTT) and insulin tolerance tests (ITT) revealed that *FicD*^*R371S/R371S*^ mice were glucose intolerant but respond well to insulin (**Fig. 1D-E**), consistent with the neonatal diabetes phenotypes described in human patients (*7*).

**Fig. 1.**
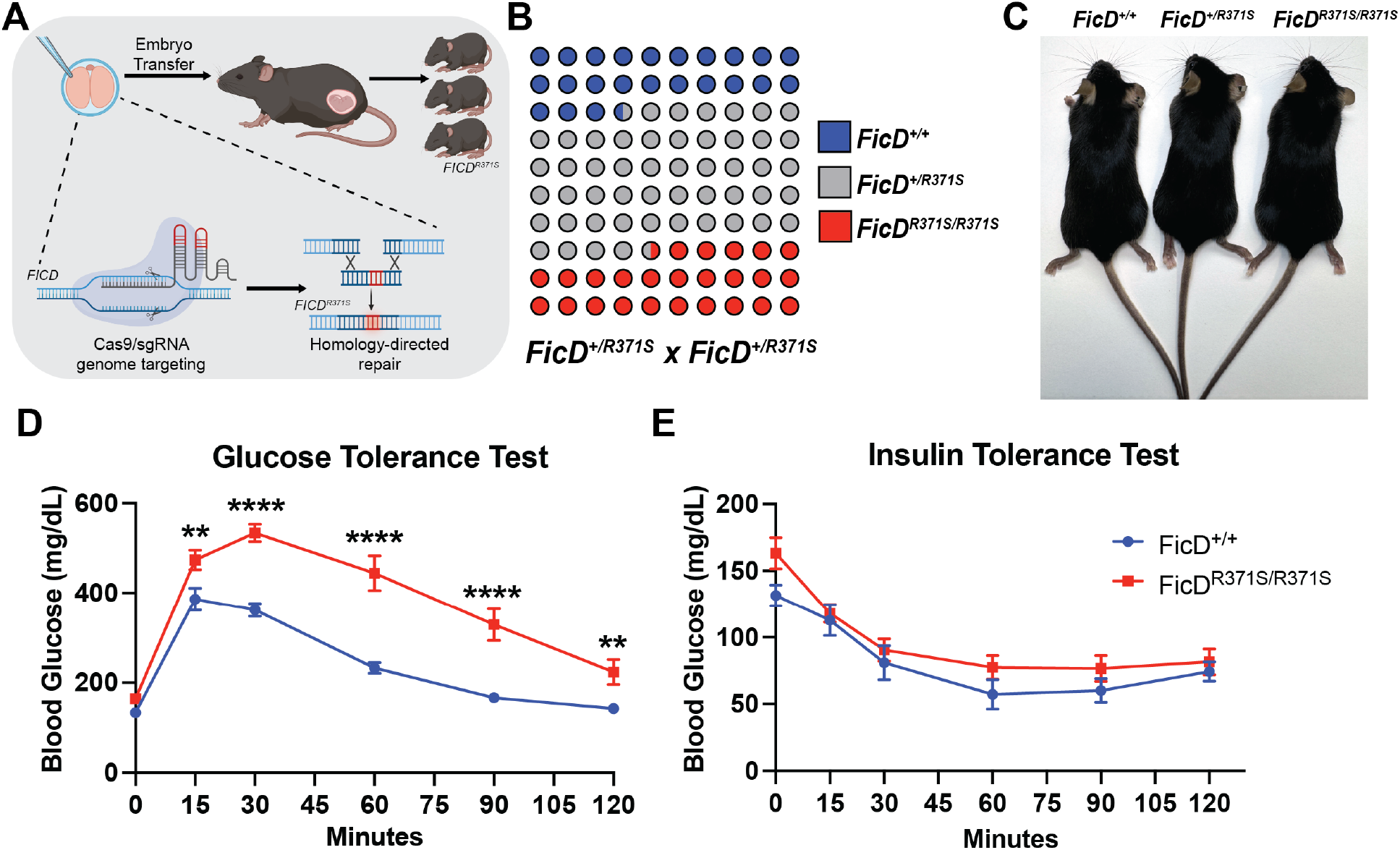
Generation of a neonatal diabetic FicD^R371S^ mouse model. (**A**) Diagram for the generation of *FicD*^*R371S*^ mice. (**B**) Schematic representing the distribution of genotypes from 14 *FicD*^*+/R371S*^ x *FicD*^*+/R371S*^ breeding cages, a total of 58 litters, 305 pups total. (**C**) Comparison of 10-week-old female *FicD*^*+/+*^, *FicD*^*+/R371S*^, and *FicD* ^*R371S /R371S*^ littermates. (**D**) GTT and (**E**) ITT of 15 to 16-week old *FicD*^*+/+*^ (blue) and *FicD*^*R371S/R371S*^ (red) mice. Data indicate mean and error bars represent standard error. Statistics were performed using GraphPad Prism 10 using an 2-way ANOVA. N= 6. **, p <0.01; ***, p <0.001; ****, p <0.0001.

### The *FicD*^*R371S*^ mutation recessively alters global AMPylation of BiP

Previous in vitro studies (*7*) have shown that FicD^R371*S*^ retains the ability to AMPylate BiP but lacks deAMPylation activity (**Fig. 2A**). Since FicD is expressed ubiquitously, we hypothesized that BiP and BiP AMPylation levels would be elevated across various tissues in *FicD*^*R371S/R371S*^ mice. Indeed, we observed significant increases in both total BiP protein and BiP AMPylation levels in multiple tissues of *FicD*^*R371S/R371S*^ mice (**Fig. 2B-D**). In contrast, AMPylation and BiP levels in *FicD*^*+/R371*S^ and FicD^+/+^ controls were indistinguishable, consistent with the recessive nature of FicD^R371S^-induced neonatal diabetes in human patients.

**Fig. 2.**
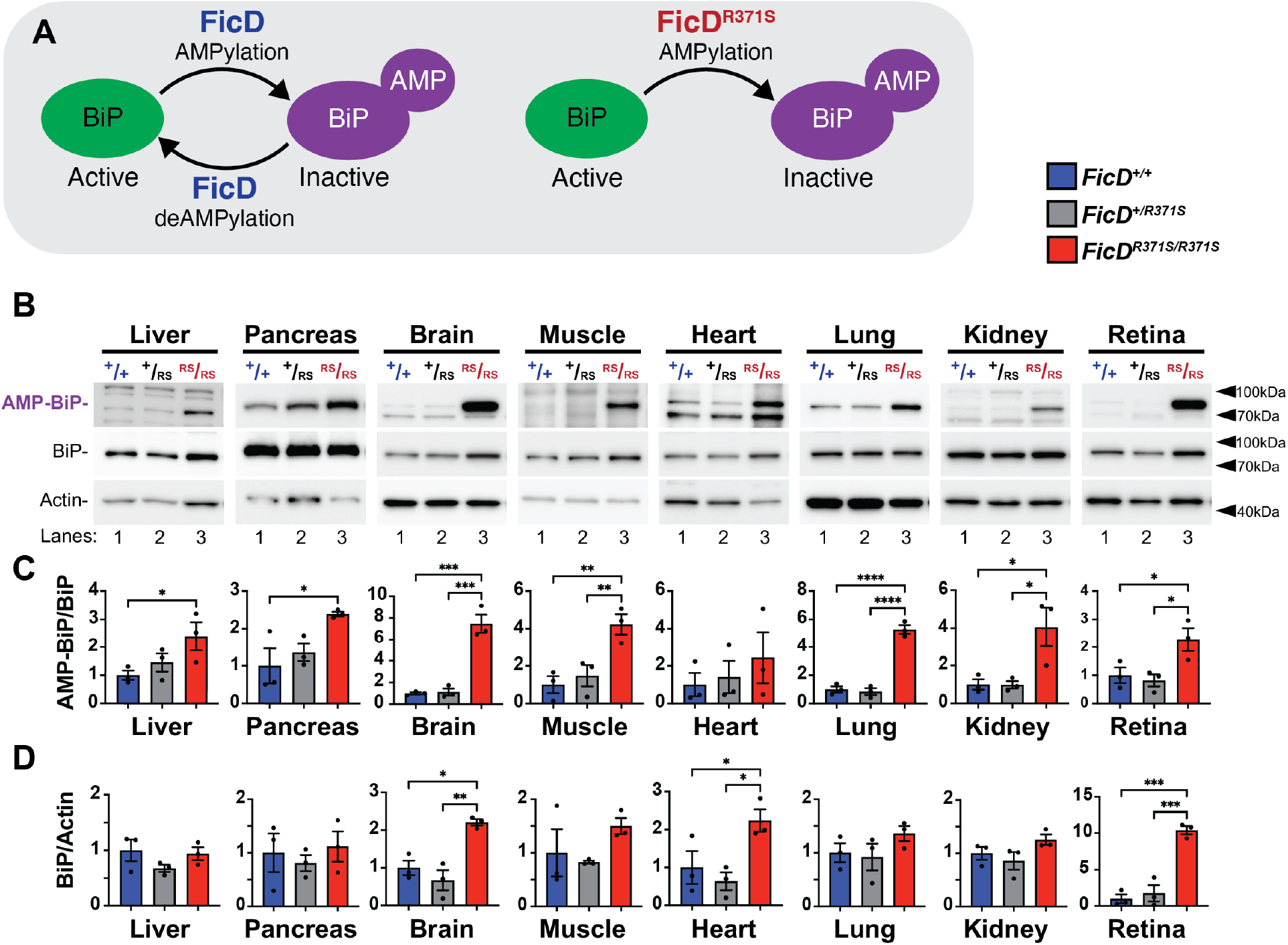
The *FicD*^*R371S*^ mutation recessively alters global AMPylation of BiP. (**A**) Model of wildtype FicD and mutant FicD^R371S^ activity. (**B**) Representative Western blots of protein lysates isolated from *FicD*^*+/+*^, *FicD*^*+/R371S*^, and *FicD* ^*R371S/R371S*^ tissues. Blots were probed with anti-AMP (17g6), anti-BiP, and anti-Actin antibodies. (**C**) Quantification of detected AMP-BiP (78kDa) relative to detected BiP. (**D**) Quantification of BiP relative to Actin. N=3. Bars indicate mean relative expression normalized to *FicD*^*+/+*^ controls, and error bars represent standard error. Statistics were performed using GraphPad Prism 10 using a 1-way ANOVA. *, p < 0.05; **, p < 0.01, ***, p < 0.001.

To test how BiP AMPylation responds to pharmacologically induced ER stress, we administered a single intraperitoneal dose of tunicamycin (Tm, 1.0mg/kg, dissolved in 150mM dextrose) and sacrificed the mice 4 hours post-injection. Control mice received vehicle injections. In response to Tm treatment, *FicD*^*+/+*^, *FicD*^*+/R371S*^, *FicD*^*R371S/R371S*^ siblings all exhibited a marked reduction in BiP AMPylation in the liver. However, in *FicD*^*R371S/R371S*^ liver, BiP AMPylation was unresponsive to tunicamycin (**Fig. S3A**). Despite these alterations in BiP AMPylation/deAMPylation dynamics under stress conditions, there were no significant differences in UPR activation across genotypes following ER stress induction (**Fig. S3B-G**).

### Loss of BiP deAMPylation alters UPR signaling in the pancreas and affects β-cell transcript expression

Given the role of FicD-mediated BiP AMPylation/deAMPylation in ER stress regulation (*8, 9*), we investigated whether UPR signaling was dysregulated in the pancreas of *FicD*^*R371S/R371S*^ mice. We found that BiP AMPylation in pancreatic lysates from *FicD*^*+/+*^ mice, but not from *FicD*^*R371S/R371S*^ mice, responded to mild physiological stress induced by fasting and refeeding (**Fig. 3 A-B**). Additionally, UPR responsive genes were transcriptionally elevated in the pancreas of *FicD*^*R371S/R371S*^ mice compared to *FicD*^*+/+*^ mice under fasting and refeeding conditions (**Fig. 3C-F**), indicating dysregulation of UPR signaling. The difference between the response to pharmacological stress (**Fig. S3A)** and physiological stress (**Fig. 3**) highlights the specific role of FicD in modulating UPR activation.

**Fig. 3.**
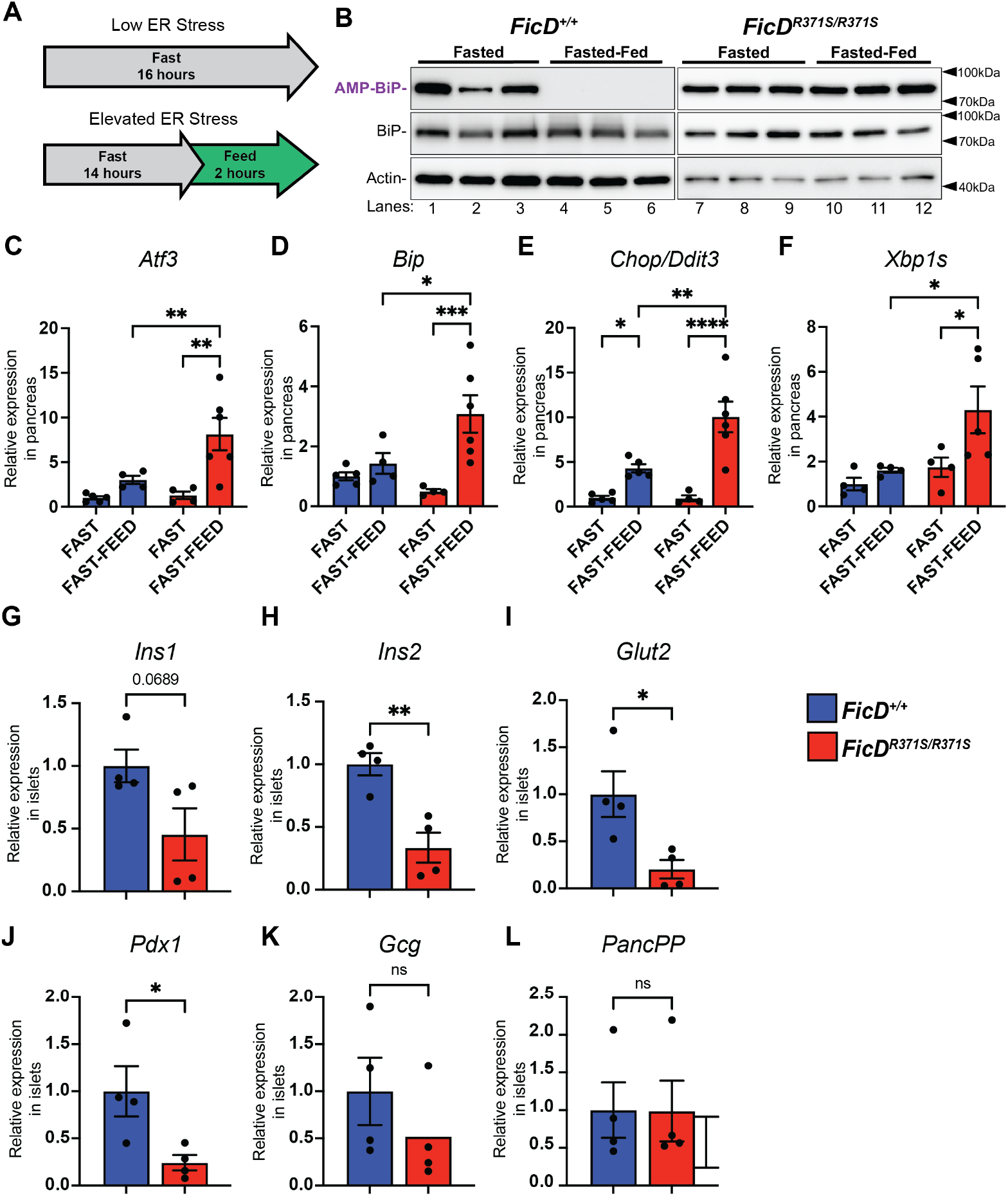
Loss of BiP deAMPylation alters UPR signaling in the pancreas and affects β-cell transcript expression. (**A**) A schematic representation of fasted, fasted-fed experimental conditions. **(B**) Representative Western blots of protein lysates isolated from *FicD*^*+/+*^ and *FicD* ^*R371S/R371S*^ pancreas. Blots were probed with anti-AMP (17g6), anti-BiP, and anti-Actin antibodies. (**C-F**) *Atf3, BiP, Chop/Ddit3*, and *Xbp1s* mRNA analyzed by qPCR from *FicD*^*+/+*^ (blue bar) and *FicD*^*R371S/R371S*^ (red bar) mouse pancreas after fasting and fast-feeding. Expression values were normalized to *Gapdh*. Bars indicate mean relative expression compared to fasted *FicD*^*+/+*^ controls, and error bars represent standard error. N=4-6. (**G-L**) *Ins1, Ins2, Glut2, Pdx1, Gcg*, and *PancPP* mRNA analyzed by qPCR from islets isolated from *FicD*^*+/+*^ (blue bar) and *FicD*^*R371S/R371S*^ (red bar) mice. Expression values were normalized to *Gapdh*. Bars indicate mean relative expression compared to *FicD*^*+/+*^ controls, and error bars represent standard error. N=4-5. Statistics were performed using GraphPad Prism 10 using (**C-F)** 2-way ANOVA or (**G-L**) unpaired student’s t-test. *, p < 0.05; **, p <0.01; ***, p <0.001; ****, p <0.0001; ns, not significant.

To examine baseline ER stress in the endocrine pancreas before fasting, we isolated pancreatic islets from *FicD*^*R371S/R371S*^ and *FicD*^*+/+*^ sibling controls and performed qPCR to assess UPR signaling- and islet function-related transcripts (**Fig. 3**). Under non-stressed conditions, UPR transcript levels in *FicD*^*R371S/R371S*^ islets were comparable to those in *FicD*^*+/+*^ controls, consistent with findings in the whole pancreas (**Fig. S4**).

However, analysis of endocrine cell-related transcripts revealed that many β-cell-specific transcripts were generally reduced in *FicD*^*R371S/R371S*^ islets (**Fig. 3G-J**). Notably, transcript levels for *Ins1* and *Ins2* (encoding insulin), *Glut2* (Glucose transporter type 2), and *Pdx1*(Pancreatic and duodenal homeobox 1) were reduced in *FicD*^*R371S/R371S*^ islets. In contrast, the α-cell (Glucagon, *Gcg*) and F-cell (Pancreatic polypeptide, *PancPP*) markers appeared unaffected.

### Regulation of FicD is required to maintain pancreatic islet organization and function

Glucose intolerance, reduced insulin secretion, and downregulation of β-cell transcripts are hallmarks of declining β-cell health and function. To assess pancreatic tissue health in *FicD*^*R371S/R371S*^ mice and their sibling controls, we performed Hematoxylin and Eosin (H&E) and TUNEL staining on paraffin embedded pancreas samples (**Fig. S5**). Histological analysis revealed no signs of inflammation or fibrosis in the islets (**Fig. S5A**), and there was no significant increase in TUNEL-positive cells, indicating a lack of significant apoptosis (**Fig. S5B**).

To further examine the organization and health of endocrine tissue, we performed immunostaining for insulin (β-cells) and glucagon (α-cells) on paraffin embedded pancreas sections from *FicD*^*+/+*^ and *FicD*^*R371S/R371S*^ siblings (**Fig. 4, Fig. S6**). Islet organization was evaluated at multiple timepoints (P0, 5 weeks, 14 weeks and 1 year) to assess potential changes in islet structure over time (**Fig 4**., **Fig. S6**).

**Fig. 4.**
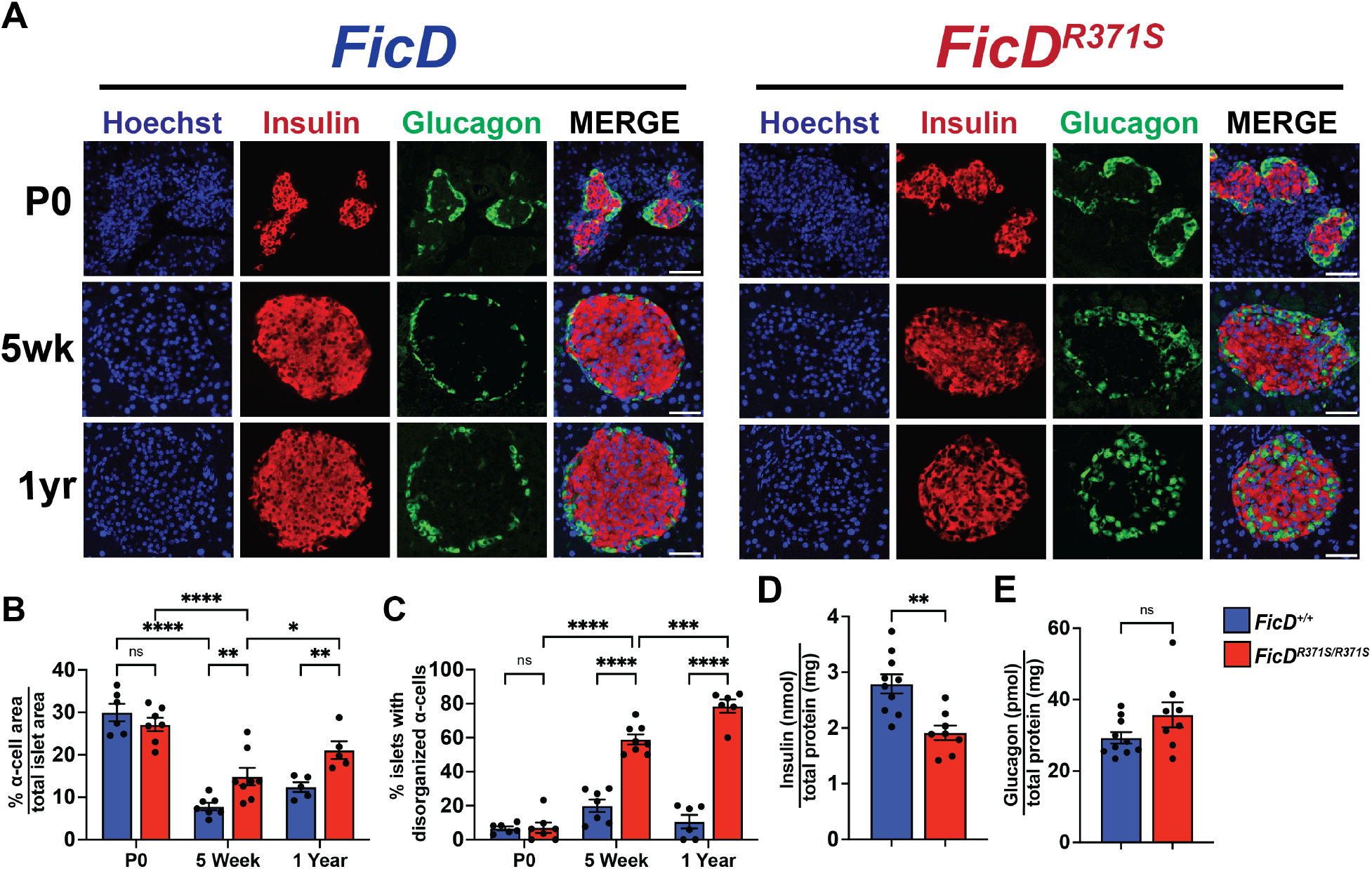
Regulation of FicD is required to maintain pancreatic islet organization and function. (**A**) Representative immunofluorescence images for insulin and glucagon expression in P0, 5-week, and 1-year-old *FicD*^*+/+*^ and *FicD*^*R371S/R371S*^ mice. Scale bar, 50uM. (**B**) Quantification of α-cells positive area as a percentage of total islet area of P0, 5-week, and 1-year-old *FicD*^*+/+*^ (blue bar) and *FicD*^*R371S/R371S*^ (red bar) mice. N=6-8. (**C**) Quantification of percent of islets with disorganized (internal to islet) α-cells of P0, 5-week, and 1-year-old *FicD*^*+/+*^ (blue bar) and *FicD*^*R371S/R371S*^ (red bar) mice. N=6-8. Bars indicate mean, and error bars represent standard error. **(D-E**) Quantification of tissue levels of insulin and glucagon from 1-year-old *FicD*^*+/+*^ (blue bar) and *FicD*^*R371S/R371S*^ (red bar) pancreas. Bars indicate mean, and error bars represent standard error. N=8-10. Statistics were performed using GraphPad Prism 10 using (**B-C**) 2-way ANOVA or (**D-E**) unpaired student’s t-test. *, p < 0.05; **, p <0.01; ***, p <0.001; ****, p <0.0001; ns, not significant.

At birth (P0), the islets in *FicD*^*R371S/R371S*^ pancreases were well organized and indistinguishable from those in *FicD*^*+/+*^ controls (**Fig. 4, Fig. S5**). However, by 5 weeks of age, *FicD*^*R371S/R371S*^ islets contained a greater number of glucagon-positive α-cells compared to controls (**Fig. 5B**). Moreover, the α-cells in *FicD*^*R371S/R371S*^ mice were more disorganized, frequently scattered throughout the islet mass (**Fig. C**). This α-cell disorganization worsened with age, as seen in 14-week-old and 1-year-old mice (**Fig. 4, Fig. S6**). Despite these changes in cell composition and organization, the size of the pancreatic islets in *FicD*^*R371S/R371S*^ mice did not significantly change with age (**Fig. S6D**)

## Discussion

Under conditions of low ER stress, FicD AMPylates BiP, thereby inactivating the chaperone and reducing its activity (**Fig. 1**). During increased ER stress, FicD switches its function to deAMPylate BIP, rapidly increasing the pool of active chaperone to handle the elevated stress (*10*). This dual catalytic activity of FicD is crucial for maintaining the appropriate level of active BiP during fluctuating ER stress conditions. In previous studies with genetic models of FicD deletion in flies and mice, we and others have observed that the absence of FicD results in tissues being less resilient to physiological and environmental stress (*8, 11, 12*).

In this study, we developed a preclinical mouse model for neonatal diabetes based on the hFicD^R371S^ mutation identified in humans. This mutation disrupts FicD’s ability to deAMPylate BiP, impairing its regulatory function. Although *FicD*^*R371S/R371S*^ mice appear overall healthy, they exhibit signs of insulin-dependent diabetes, including reduced glucose tolerance and diminished insulin secretion. While the islets of *FicD*^*R371S/R371S*^ mice are normally organized at birth, they become progressively disorganized with age. Furthermore, we observed altered β-cell transcript expression in isolated *FicD*^*R371S/R371S*^ islets and elevated UPR signaling in the pancreas during fasting-feeding stress.

Taken together, our data suggest that the pancreatic defects in *FicD*^*R371S/R37S*^ mice are progressive, likely resulting from cumulative disfunction due to repeated islet stress. Since the pancreas appears to develop normally in utero, when blood glucose levels are regulated by the mother, we postulate that the postnatal defects in the *FicD*^*R371S/R37S*^ mice arise over time as a response to physiological stress, particularly related to regulating blood glucose levels after birth.

In humans, patients homozygous for *hFicD*^*R371S*^ mutation present with symptoms across multiple organs, including infancy-onset diabetes, severe neurodevelopmental delays, short stature, and cataracts. Similarly, we observed elevated BiP AMPylation levels across multiple tissues in *FicD*^*R371S/R371S*^ mice, suggesting that BiP function is globally altered. Future studies are warranted to assess the molecular consequence of FicD dysregulation in other secretory tissues affected by *hFicD*^*R371S*^, to better understand how this mutation impacts tissue health and UPR signaling under chronic physiological stress.

## Acknowledgments

We thank Phillip Scherer and Shiuhwei Chen for their valuable discussions and advice. We thank UT Southwestern Metabolic Phenotyping Core for the analysis of GTT and ITT and expertise. We thank Aymelt Itzen for his generous gift of monoclonal anti-adenosine monophosphate antibodies. Fig. 1A prepared with source icons created with BioRender.com

## Funding

National Institutes of Health Grant R35 GM134945 (KO) National Institutes of Health Grant R01 HD103627-01A1 (JW)

National Institutes of Health Grant R21 1R21EY034597-01A1 (AKC) New York Stem Cell Foundation GM138565-01A1 (JW)

New York Stem Cell Foundation–Robertson Investigator and Virginia Murchison Linthicum Scholar in Medical Research (JW)

Once Upon a Time…Foundation (KO)

UTSW-NORC grant under National Institute of Diabetes and Digestive and Kidney Disease/National Institutes of Health award number P30DK127984 (AKC).

Welch Foundation grant I-1561 (KO)

WW Caruth, Jr. Biomedical Scholar with an Earl A Forsythe Chair in Biomedical Science (KO)

## Author contributions

Conceptualization: AKC, JW, KO

Investigation: AKC, NMS, CAP, MS, NZ, HFG, HAF

Visualization: AKC, NMS

Funding acquisition: AKC, KO Project administration: AKC, KO

Supervision: AKC, KO

Writing – original draft: AKC, KO

Writing – review & editing: AKC, KO, JW, BME,

## Data and materials availability

All data are available in the main text or the supplementary materials.

## Supplementary Materials

Materials and Methods Figs. S1 to S6

Table S1

