## Supplementary material for "Dysregulation of FicD AMPylation causes diabetes by disrupting pancreatic endocrine homeostasis": Supp info

**The PDF file includes:**

Materials and Methods
Figs. S1 to S6
Tables S1

### Materials and Methods

Mice

The Institutional Animal Care and Use Committee of the University of Texas Southwestern Medical Center approved all experiments. C57BL/6J and BDF1 mice were purchased from The Jackson Laboratory. CD-1 (ICR) mice were purchased from Envigo (Harlen). Mice were housed in a specific pathogen-free facility with a 12 hour light dark cycle and fed an *ad libitum* standard chow diet (#2016 Harlan Teklad), unless specified. Fasting and fasting feeding studies were performed as previously described with 6-8 week old males (*8*). Fasting was performed overnight, with 2 hour feeding occurring at 8am. Administration of 1.0mg/kg of tunicamycin was performed as previously described (*13*). Briefly, 8-10 week old male mice were administered a single intraperitoneal dose (1.0mg/kg) of tunicamycin (Sigma) dissolved in 150mM dextrose at approximately 10am and sacrificed after 4 hours. Mice in the control group were injected with vehicle alone.

Cas9 mRNA in vitro transcription

We modified the PX458 plasmid (Addgene plasmid # 48138) by adding T7 promoter upstream of Cas9 coding sequence and removing T2A-GFP. The modified plasmid was linearized by Not I (NEB) digestion. The linearized Cas9 plasmid and PCR products were purified using QIAquick PCR Purification Kit (QIAGEN). Cas9 mRNA was in vitro transcribed using purified linearized plasmid as a template by mMESSAGE mMACHINE™ T7 Transcription Kit (Invitrogen). Prepared Cas9 mRNA was then purified by Lithium chloride precipitation and dissolved in water for embryo transfer (Sigma: W1503).

Design sgRNA and donor repair template

The single guide RNAs (sgRNAs) and single-stranded repair template were design as described in (*14*). Briefly DNA sequences where imported intro benching (https://benchling.com, 2022), and sgRNAs that cut near (<10bp) to the intended mutation site were designed with the Benchling design tool. sgRNAs were filtered by the folding quality using the WU-CRISPR server (https://crisprdb.org/wu-crispr-website/). The gRNA sequence (5'CTTTCATCGACGGCAACGG'3) was predicted to cut approximately 3 bp from our targeted mutation site and sgRNA was purchased from Synthego. The single-stranded oligodeoxynucleotide (ssODN) repair template, complementary to the PAM distal strand with 5' and 3' phosphorothioate ends to prevent exonuclease degradation and enhance HDR repair efficiency was purchased from IDT. (5'gaggacgccatgaacctgcacccagtcgagttcgcggccctggcccattacaaactggtgtacatccaccctttcatcgacggcaacg gATCCacctcccgtctgctgatgaacctgattttgatgcaggcgggataccc'3), were the R370S mutation disrupts the PAM sequence and contains a silent mutation in G369 (GGG>GGA) to create a new BamHI site.

Embryo collection

Harvesting embryos was performed as described previously with slight modification (*15, 16*). Briefly, BDF1 female mice (4 weeks) were superovulated by intraperitoneal (IP) injection with 7.5 IU of PMSG (Prospec: #HOR-272), followed by IP injection of 7.5 IU of hCG (Sigma: #C1063) 48 later. After mating with C57BL/6J male mice, zygotes were harvested at E0.5 (the presence of a virginal plug was defined as embryonic day 0.5 (E0.5)) in mKSOM-Hepes from oviducts and cumulus cells were removed with hyaluronidase (Sigma: #H4272). Zygotes were cultured in the mKSOMaa until Cas9 mRNA/sgRNA/donor ssODN microinjection in a humidified atmosphere containing 5% (v/v) $CO_2$ at 37 °C.

Microinjection of Cas9 mixture to zygotes

Microinjection of Cas9 mRNA/sgRNA/donor ssODN was performed as described previously (*15*) with slight modification. Briefly, the zygotes show clear two pronuclei were selected and transferred into a 40 µL drop of KSOM-Hepes and placed on an inverted microscope (Nikon, Japan) fitted with micromanipulators (Narishige, Japan). Mixture of Cas9 mRNA (100 ng/µL), sgRNA (50 ng/µL) and donor ssODN (20 ng/µL) was loaded to a blunt-end micropipette (Sutter Instrument, CA) of 2–3 µm internal diameter, and Piezo Micro Manipulator (Prime Tech Ltd, Japan) was used to create a hole in the zona pellucida and the zygote membranes. The injection of the Cas9 mixture into the cytoplasm of zygotes was confirmed by the bulge of membrane. Groups of 12 zygotes were manipulated simultaneously and each session was limited to 10 min. After microinjection, the zygotes were cultured in the 40 µL droplet of mKSOMaa for 3 days in a humidified atmosphere of 5% $CO_2$, 5% $O_2$ in air at 37 °C.

Embryo transfer

Embryo transfer was performed as described previously(*15, 16*) . Briefly, CD-1 female mice (8 weeks old or older) were mated with vasectomized CD-1 male mice to induce pseudopregnancy. 8–13 embryos at E3.5 were surgically transferred to the surrogate uterine at E2.5 under anesthesia with Ketamine (30 mg/mL)/Xylazine (4 mg/mL) and analgesia with Buprenorphine SR-LAB (1 mg/mL) within 20–30 min per surrogate. F0 mice were bred with C57Bl/6J mice to obtain F1 mice heterozygous for the $FicD^{R371S}$ mutation.

Genotyping and DNA sequencing

To determine genotypes of full-term delivered pups, tail-tips were used for genomic DNA extraction using DNeasy Blood and Tissue kit (Qiagen). The genomic DNA sequences including target site were amplified with PrimeSTAR GXL DNA Polymerase. PCR products were purified with QIAquick PCR Purification kit (Qiagen) and submitted for Sanger sequencing (Eurofins Genomics) to confirm the $FicD^{R371S}$ mutation. Forward primer: 5' tcctgcacgccctcaagatgga 3' Reverse primer: 5' cgtcaccctcgttggcgacttc 3'. All mice used in this study were genotyped via PCR using the primer set (5' gggggtggttcaaggaag 3' and downstream 5' ctgcaacctcctactggc 3' followed by restriction digestion with BamH1). Amplicons of total FicD gene were submitted for Sanger sequencing to confirm the $FicD^{R371S}$ mutation. F0 mice were bred with C57Bl/6J mice to obtain F1 mice heterozygous for the $FicD^{R371S}$ mutation.

Glucose Tolerance Test and Insulin Tolerance Test

For glucose tolerance tests (GTTs) and insulin tolerance tests (ITTs), mice were fasted for 6 hours with water provided ad libitum starting at 7 to 8 am on the experimental day. Baseline glucose levels were collected 1 hour prior to GTT or ITT. During GTT or ITT, blood glucose levels were monitored at 0, 15, 30, 60, 90, and 120 minutes after an i.p. 2.0 g/kg body weight dose of glucose (Gibco 20% ref A24940-01, lot 2562566) or 1U/kg body weight dose of Insulin (Humalin), respectively. Blood samples were collected from the tail vein and glucose was measured using a glucometer.

Islet Isolations

Pancreatic islets were isolated from mice as previously described (*17*). Briefly, the pancreas was digested via injection through the pancreatic duct with Collagenase in Hank's balanced salt solution. Digestion was halted through washes with Hank's balanced salt solution and islet/exocrine tissue was transferred to RPMI 1640. Islets were hand-picked before used in experiments.

Assessment of Serum Pancreatic Insulin and Glucagon Levels.

Insulin and glucagon were extracted from the pancreas using the acid ethanol method(*18*). For serum isolation, blood was collected in Microvette serum collection tubes (Sarstedt) and centrifuged at 10,000xg at 4°C for 15 minutes. Insulin and Glucagon levels were measured using a commercially available ELISA kits (Crystal Chem., Inc.).

Histology and Immunohistochemistry

Mouse pancreas were harvested and fixed in 10% neutral buffered formalin overnight at 4°C. Paraffin sections were embedded by the UT Southwestern (UTSW) Histo Pathology Core. Hematoxylin and eosin (H&E) staining was performed using standard techniques(*19*). Nuclei were visualized with Hoechst. TUNEL/PI staining was performed by the UTSW Histo Pathology Core using the DeadEnd™ Fluorometric TUNEL system (Promega). Pancreatic glucagon and insulin distribution was determined using deparaffinized pancreas section after a hot citric acid buffer (10mM citric acid, 0.05% Tween-20, pH 6.0) antigen retrieval. Sections were incubated with anti-insulin (Abcam ab181547) and anti-glucagon (Invitrogen 14-9743-82) antibodies at 4°C for 16 hours. Secondary antibodies used were Alexa Fluor 488 goat anti-mouse (Invitrogen) and Alexa Fluor 555 goat anti-rabbit (Invitrogen)..

Quantitative real-time PCR

RNA from total pancreas and isolated islets was extracted using RNA Stat-60 (Iso-Tex Diagnostics). Complementary DNA (cDNA) was generated from RNA (2 μg) using the High-Capacity cDNA Reverse Transcription Kit (Life Technologies). qPCR was performed by the SYBR Green method(*20*). DNA contamination in RNA and reagents was assessed using no reverse transcriptase samples in which qPCR reactions were performed on mock cDNA from samples prepared without reverse transcriptase and no template control samples in which no template was added to the qPCR reaction. Primer sequences for the genes analyzed can be found in **Table S1**. Experiments were performed on a BioRad CFX Touch and analyzed with CFX Maestro Software.

Western Blot Analysis

Collected tissues were washed in PBS and homogenized in RIPA buffer (50mM Tris pH 8, 150mM NaCl, 1% NP-40, 0.5% Na deoxycholate, PMSF, PhosSTOP (Roche), Protease Inhibitor Cocktail (Roche. Lysates were centrifuged at 10,000xg for 10 minutes to remove nuclei and cellular debris. Lysates were separated by SDS-PAGE and transferred to PVDF membranes. Blots were probed with anti-AMP 17g6 (gift from Aymelt Itzen) (*21*), anti-GRP78 (Abcam, ab21685), and anti-actin (Sigma A2228). Membranes were then incubated with horseradish peroxidase–conjugated secondary antibodies Goat Anti-Mouse IgG-HRP (Abcam, ab205719) or Donkey anti-

Rabbit IgG-HRP (Fisher, NA934) against the primary antibody's host species for 1 hour. Membranes were developed using the ECL substrate solution (Bio-Rad). Quantification of western blots was performed using NIH ImageJ software. Band densities were measured and subtracted from background.

**A**

```
FicD       1021  GAGGACGCCATGAACCTGCACCCAGTCGAGTTCGCGGCCCTGGCCCATTACAAACTGGTG  1080
                 ||||||||||||||||||||||||||||||||||||||||||||||||||||||||||||
FicDR371S  1021  GAGGACGCCATGAACCTGCACCCAGTCGAGTTCGCGGCCCTGGCCCATTACAAACTGGTG  1080

FicD       1081  TACATCCACCCTTTCATCGACGGCAACGGGAGGACCTCCCGTCTGCTGATGAACCTGATT  1140
                 ||||||||||||||||||||||||||||||    ||||||||||||||||||||||||||
FicDR371S  1081  TACATCCACCCTTTCATCGACGGCAACGGATCCACCTCCCGTCTGCTGATGAACCTGATT  1140

FicD       1141  TTGATGCAGGCGGGATACCCGCCCATCACCATCCGCAAGGAGCAGAGGTCCGAGTACTAC  1200
                 ||||||||||||||||||||||||||||||||||||||||||||||||||||||||||||
FicDR371S  1141  TTGATGCAGGCGGGATACCCGCCCATCACCATCCGCAAGGAGCAGAGGTCCGAGTACTAC  1200
```

**B**

```
FicD       1    MILMPMASVVAVAEPKWVSVWGRFLWMALLSMALGSLLALLLPLGVVEEHCLAVLRGFHL  60
                ||||||||||||||||||||||||||||||||||||||||||||||||||||||||||||
FicDR371S  1    MILMPMASVVAVAEPKWVSVWGRFLWMALLSMALGSLLALLLPLGVVEEHCLAVLRGFHL  60

FicD       61   LRSKLDRAQPVVPKCTSLCTELSVSSRDAGLLTVKTTASPAGKLEAKAALNQALEMKRQG  120
                ||||||||||||||||||||||||||||||||||||||||||||||||||||||||||||
FicDR371S  61   LRSKLDRAQPVVPKCTSLCTELSVSSRDAGLLTVKTTASPAGKLEAKAALNQALEMKRQG  120

FicD       121  KRGKAHKLFLHALKMDPGFVDALNEFGIFSEEDKDIIQADYLYTRALTISPFHEKALVNR  180
                ||||||||||||||||||||||||||||||||||||||||||||||||||||||||||||
FicDR371S  121  KRGKAHKLFLHALKMDPGFVDALNEFGIFSEEDKDIIQADYLYTRALTISPFHEKALVNR  180

FicD       181  DRTLPLVEEIDQRYFSVIDSKVKKVMSIPKGSSALRRVMEETYYHHIYHTVAIEGNTLTL  240
                ||||||||||||||||||||||||||||||||||||||||||||||||||||||||||||
FicDR371S  181  DRTLPLVEEIDQRYFSVIDSKVKKVMSIPKGSSALRRVMEETYYHHIYHTVAIEGNTLTL  240

FicD       241  SEIRHILETRYAVPGKSLEEQNEVIGMHAAMKYINTTLVSRIGSVTMDDMLEIHRRVLGY  300
                ||||||||||||||||||||||||||||||||||||||||||||||||||||||||||||
FicDR371S  241  SEIRHILETRYAVPGKSLEEQNEVIGMHAAMKYINTTLVSRIGSVTMDDMLEIHRRVLGY  300

FicD       301  VDPVEAGRFRRTQVLVGHHIPPHPRDVEKQMQEFTQWLNSEDAMNLHPVEFAALAHYKLV  360
                ||||||||||||||||||||||||||||||||||||||||||||||||||||||||||||
FicDR371S  301  VDPVEAGRFRRTQVLVGHHIPPHPRDVEKQMQEFTQWLNSEDAMNLHPVEFAALAHYKLV  360

FicD       361  YIHPFIDGNGRTSRLLMNLILMQAGYPPITIRKEQRSEYYHVLEVANEGDVRPFIRFIAK  420
                |||||||||| ||||||||||||||||||||||||||||||||||||||||||||||||
FicDR371S  361  YIHPFIDGNGSTSRLLMNLILMQAGYPPITIRKEQRSEYYHVLEVANEGDVRPFIRFIAK  420

FicD       421  CTEVTLDTLLLATTEYSVALPEAQPNHSGFKETLPVRP  458
                ||||||||||||||||||||||||||||||||||||||
FicDR371S  421  CTEVTLDTLLLATTEYSVALPEAQPNHSGFKETLPVRP  458
```

**C**

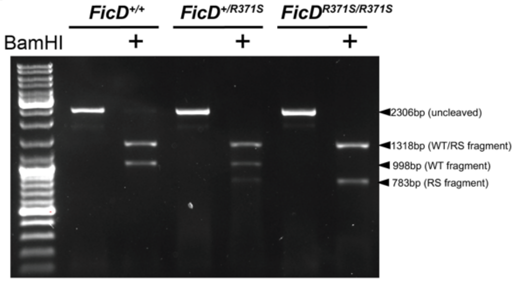


**Fig. S1.** (**A**) The aligned predicted DNA sequences of *FicD and FicD*$^{R371S}$ from nt1021-1200 . FicD is identical to Fic$^{R371S}$ except for 4 nucleotide mutations that confer a single amino acid change and silent BamHI restriction site. (**B**) The aligned predicted protein sequences of FicD and FicD$^{R371S}$. FicD is identical to Fic$^{R371S}$ except for R to S point mutation at residue 371. (**C**) Representative agarose gel of PCR amplicons and digests for FicD genotyping. Primers were used that amply across the modified region of *FicD*, producing a 2306bp *FicD* amplicon. Digestion of *FicD* amplicons with BamHI result in DNA fragments of 1318 and 988 bp (FicD), 1318 and 783 bp (FicD$^{R371S}$).

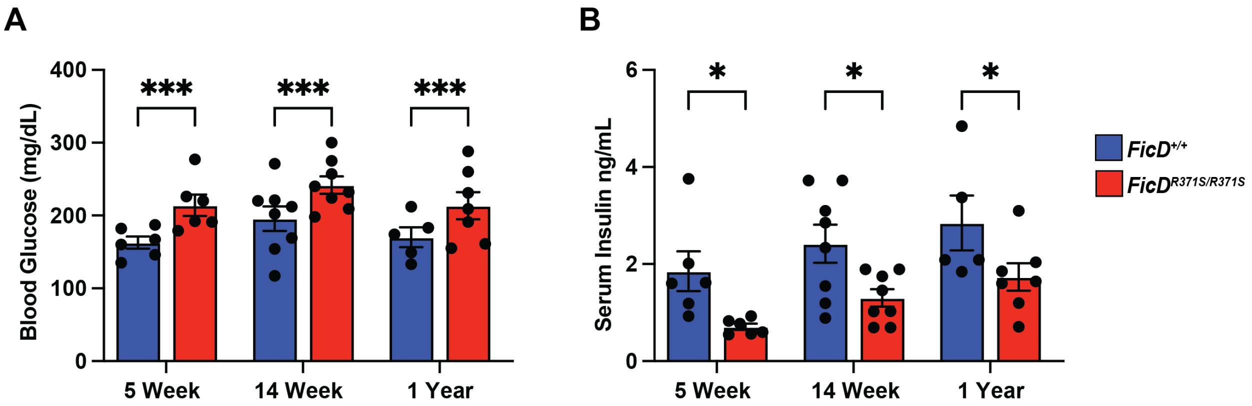


**Fig. S2.** Quantification of (**A**) blood glucose levels and (**B**) serum insulin levels of 5-week, 14-week, and 1-year-old *FicD*$^{+/+}$ (blue bar) and *FicD*$^{R371S/R371S}$ (red bar) mice. N=6-8. Bars indicate mean, and error bars represent standard error. Statistics were performed using GraphPad Prism 10 using 2-way ANOVA. *, $p < 0.05$; *** , $p < 0.001$.

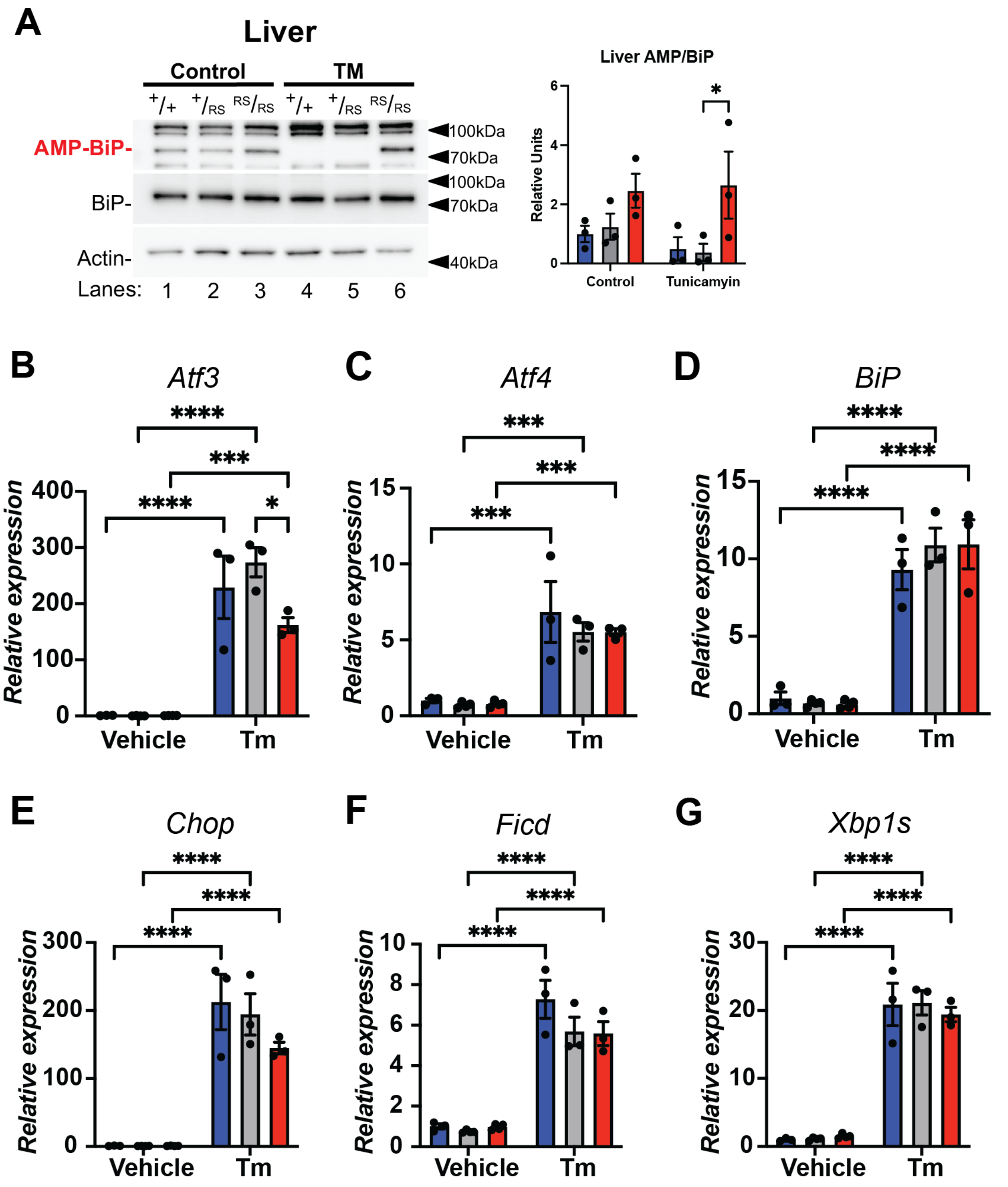


**Fig. S3.** (**A**) Representative Western blots of protein lysates isolated from *FicD*$^{+/+}$, *FicD*$^{+/R371S}$, and *FicD*$^{R371S/R371S}$ liver treated with 1mg/kg Tunicamycin or control. Blots were probed with anti-AMP (17g6), anti-BiP, and anti-Actin antibodies. Quantification of detected AMP-BiP relative to detected BiP and normalized to *FicD*$^{+/+}$ control. (**B-G**) *Atf3*, *Atf4*, *BiP*, *Chop/Ddit3*, *Ficd*, and *Xbp1s* mRNA analyzed by qPCR from *FicD*$^{+/+}$ (blue bar), *FicD*$^{+/R371S}$ (gray bar), and *FicD*$^{R371S/R371S}$ (red bar) mouse pancreas after 4hrs administration of vehicle control or 1mg/kg Tunicamycin (Tm). Expression values were normalized to *Gapdh*. Bars indicate mean relative expression compared to vehicle *FicD*$^{+/+}$ controls, and error bars represent standard error. N=3. Statistics were performed using GraphPad Prism 10 using 2-way ANOVA *, $p < 0.05$; ** , $p < 0.01$; *** , $p < 0.001$; **** , $p < 0.0001$; ns, not significant.

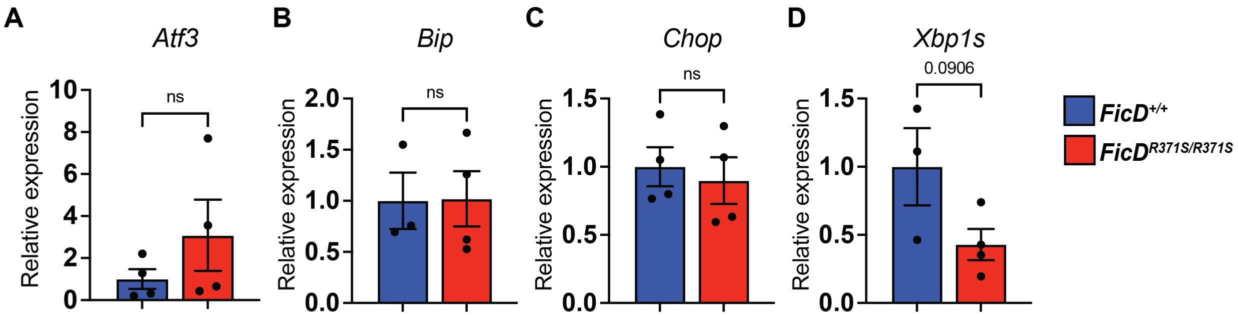


**Fig. S4.** (**A-D**) *Atf3*, *BiP*, *Chop/Ddit3*, and *Xbp1s* mRNA analyzed by qPCR islets isolated from *FicD*$^{+/+}$ (blue bar) and *FicD*$^{R371S/R371S}$ (red bar) mice. Expression values were normalized to *Gapdh*. Bars indicate mean relative expression compared to *FicD*$^{+/+}$ controls, and error bars represent standard error. N=4-5. Statistics were performed using GraphPad Prism 10 using unpaired student's t-test. ns, not significant.

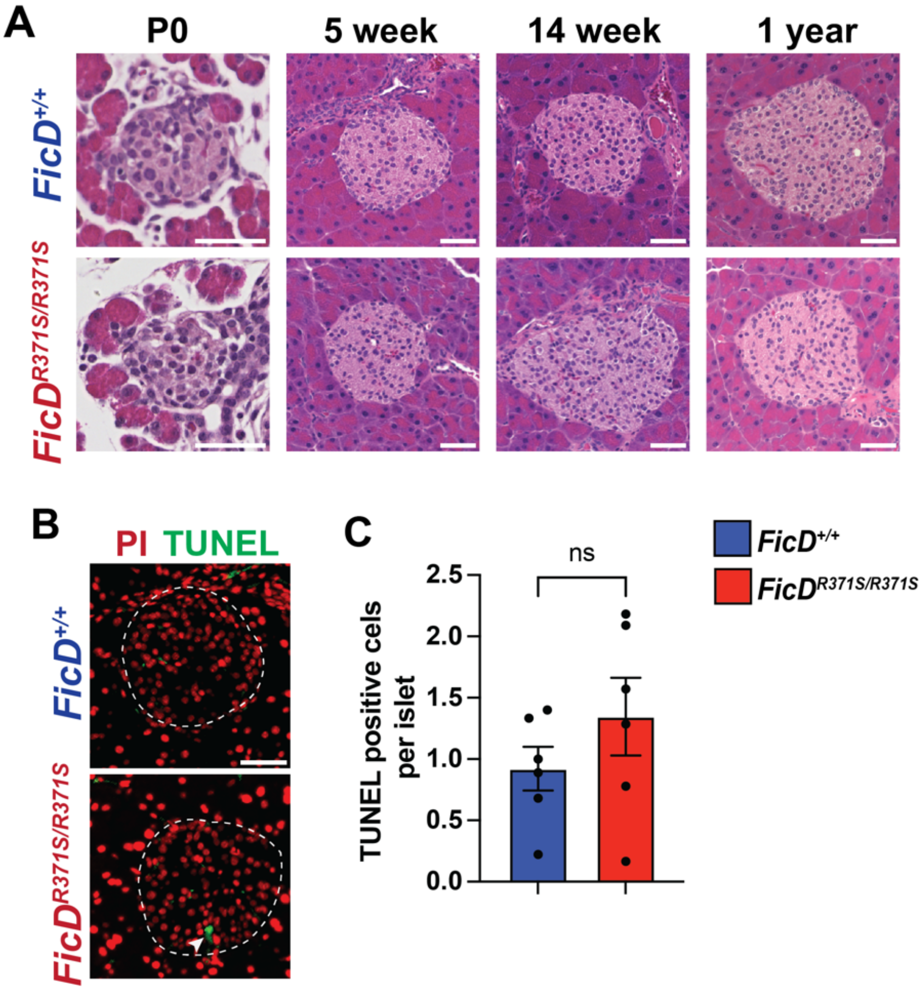


**Fig. S5.** (**A**) Representative H&E (hematoxylin and eosin) in P0, 5-week, 14-week, and 1-year-old *FicD*$^{+/+}$ and *FicD*$^{R371S/R371S}$ mice. (**B**) Representative TUNEL (Terminal deoxynucleotidyl transferase dUTP nick end labeling) and PI (propidium iodide) stained images of pancreas of 5 week old *FicD*$^{+/+}$(blue bar) and *FicD*$^{R371S/R371S}$ (red bar) mice. Scale bar, 50uM. (**C**) Quantification of percent TUNEL positive cells per islet of 5 week old *FicD*$^{+/+}$ and *FicD*$^{R371S/R371S}$ mice. Bars indicate mean, and error bars represent standard error. N=6. Statistics were performed using GraphPad Prism 10 using unpaired student's t-test. ns, not significant.

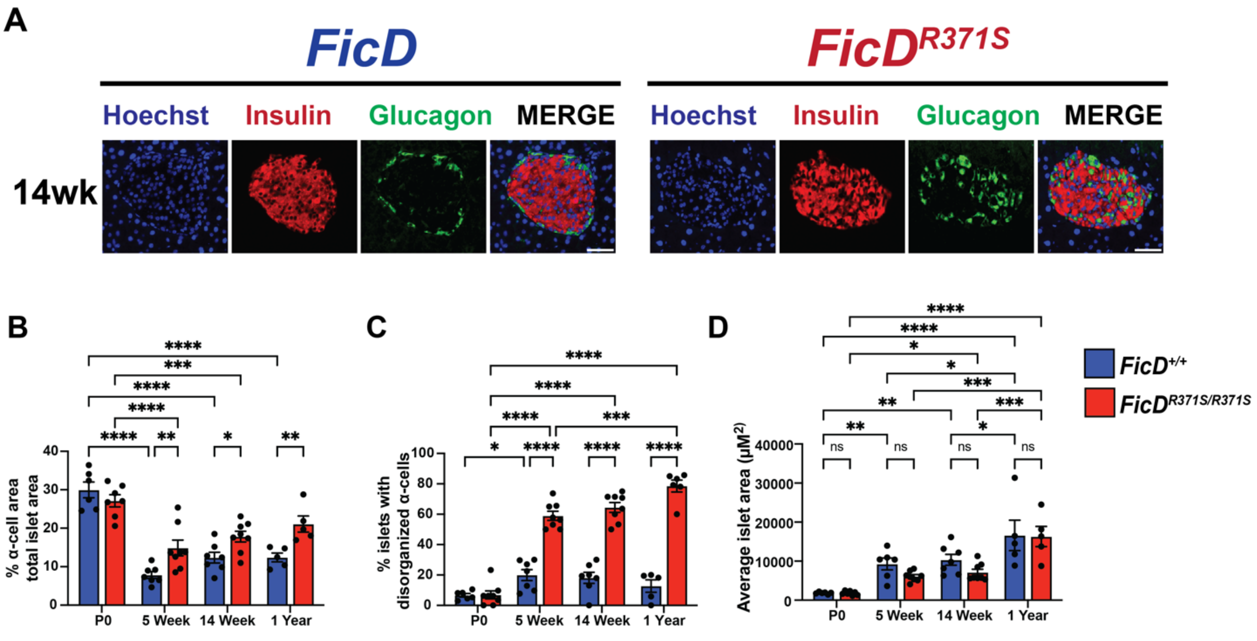


**Fig. S6.** (**A**) Representative immunofluorescence images for insulin and glucagon expression in 14 week old *FicD*$^{+/+}$ and *FicD*$^{R371S/R371S}$ mice. Scale bar, 50uM. (**B**) Quantification of α-cells positive area as a percentage of total islet area of P0, 5-week, 14-week, and 1-year-old *FicD*$^{+/+}$ (blue bar) and *FicD*$^{R371S/R371S}$ (red bar) mice. (**C**) Quantification of percent of islets with disorganized (internal to islet) α-cells of P0, 5-week, 14-week, and 1-year-old *FicD*$^{+/+}$ (blue bar) and *FicD*$^{R371S/R371S}$ (red bar) mice. (**D**) Quantification of islet area of P0, 5-week, 14-week, and 1-year-old *FicD*$^{+/+}$ (blue bar) and *FicD*$^{R371S/R371S}$ (red bar) mice. Bars indicate mean, and error bars represent standard error. N=6-8. Statistics were performed using GraphPad Prism 10 using (**B-C**) 2-way ANOVA or (**D-E**) unpaired student's t-test. *, $p < 0.05$; ** , $p <0.01$; *** , $p <0.001$; **** , $p <0.0001$; ns, not significant.

**Table S1.**

Primer sequences for the genes analyzed in this study by qPCR

| Gene name | Forward primer sequence | Reverse primer sequence |
|---|---|---|
| *Atf3* | 5’ TGGAGATGTCAGTCACCAAGTCT 3’ | 5’ GCAGCAGCAATTTTATTTCTTTCT 3’ |
| *Atf4* | 5’ ACTCTAATCCCTCCATGTGTAAAGG 3’ | 5’ CAGGTAGGACTCTGGGCTCAT 3’ |
| *BiP* | 5’ CAAGGATTGAAATTGAGTCCTTCTT 3’ | 5’ GGTCCATGTTCAGCTCTTCAAA 3’ |
| *Chop/ Ddit3* | 5’ CCAGAAGGAAGTGCATCTTCA 3’ | 5’ ACTGCACGTGGACCAGGTT 3’ |
| *Ficd* | 5’ GTAGACGCACTGAATGAGTTCG 3’ | 5’ TGGTGTATAAGTAGTCAGCCTGG 3’ |
| *Gapdh* | 5’ AGGTCGGTGTGAACGGATTTG 3’ | 5’ TGTAGACCATGTAGTTGAGGTCA 3’ |
| *Xbp1S* | 5’CTGAGTCCGCAGCAGGT 3’ | 5’ TGTCAGAGTCCATGGGAAGA 3’ |
| *Ins1* | 5’ GCCATGTTGAAACAATGACCT 3’ | 5’ CAGAGAGGAAGGTACTTTGGACTATAA 3’ |
| *Ins2* | 5’ GAAGTGGAGGACCCACAAGT 3’ | 5’ AGTGCCAAGGTCTGAAGGTC 3’ |
| *Glut2* | 5’ ACACCGGAATGTTCTTAGCC 3’ | 5’ GTGAGAAGCCGAGGAAAG 3’ |
| *Pdx1* | 5’ GAAATCCACCAAAGCTCACG 3’ | 5’ CGGGTTCCGCTGTGTAAG 3’ |
| *Gcg* | 5’ GCCCTTCAAGACACAGAGGA 3’ | 5’ CCTCATGCGCTTCTGTCTGv |
| *PancPP* | 5’ TCACTAGCTCAGCACACAGGA 3’ | 5’ CCACCCAAGTGGATACGAGA 3’ |
